# Microbial Changes occurring during oronasal fistula wound healing

**DOI:** 10.1101/2023.06.02.543508

**Authors:** Steven L. Goudy, Heath Bradley, Camillo Anthony Gacasan, Afra Toma, Crystal R. Naudin, William M. Wuest, Martin Tomov, Vahid Serpooshan, Ahmet Coskun, Rheinallt M. Jones

**Author notes:** **Corresponding author:** Steven L. Goudy, Division of Otolaryngology, Department of Pediatrics, Emory University School of Medicine, Atlanta, GA, 30322.

## Abstract

The oral microbiome is a complex community that matures with dental development while oral health is also a recognized risk factor for systemic disease. Despite the oral cavity having a substantial microbial burden, healing of superficial oral wounds occurs quickly and with little scarring. By contrast, creation of an oro-nasal fistula (ONF), often occurring after surgery to correct a cleft palate, is a significant wound healing challenge that is further complicated by a connection of the oral and nasal microbiome. In this study, we characterized the changes in the oral microbiome of mice following a freshly inflicted wound in the oral palate that results in an open and unhealed ONF. Creation of an ONF in mice significantly lowered oral microbiome alpha diversity, with concurrent blooms of *Enterococcus faecalis, Staphylococcus lentus, and Staphylococcus xylosus* in the oral cavity. Treatment of mice with oral antibiotics one week prior to ONF infliction resulted in a reduction in the alpha diversity, prevented *E. faecalis* and *S. lentus, and S. xylosus* blooms, but did not impact ONF healing. Strikingly, delivery of the beneficial microbe *Lactococcus lactis* subsp. cremoris (LLC) to the wound bed of the freshly inflicted ONF via a PEG-MAL hydrogel vehicle resulted in rapid healing of the ONF. Healing of the ONF was associated with the maintenance of relatively high microbiome alpha diversity, and limited the abundance of *E. faecalis* and *S. lentus, and S. xylosus* in the oral cavity. These data demonstrate that a freshly inflicted ONF in the murine palate is associated with a dysbiotic oral microbiome state that may prevent ONF healing, and a bloom of opportunistic pathogens. The data also demonstrate that delivery of a specific beneficial microbe, LLC, to the ONF can boost wound healing, can restore and/or preserve oral microbiome diversity, and inhibit blooms of opportunistic pathogens.

## Introduction

The oral cavity has the second most diverse microbiota after the gut with the community structure totaling over 700 species of bacteria. The oral microbiome is crucial in maintaining oral, as well as systemic health. Initial colonizers immediately after birth include mainly aerobes such as *Streptococcus, Lactobacillus, Actinomyces, Neisseria* and *Veillonella*. Thereafter, following tooth eruption and plaque accumulation, and the development of gingival crevices, wider species diversity and microbial succession develops within the oral cavity until microbes from the phyla *Firmicutes, Proteobacteria, Bacteroidetes, Actinobacteria, Fusobacteria, Neisseria*, predominately inhabit the oral cavity. The fundamentals of oral cavity microbiome have been outlined in several review articles^1-8^. However, less is known about the oral microbiome in relation to large mucosal injury as would occur following cleft palate corrective surgery, although a recent report on the oral microbiome of children with cleft lip and palate revealed associations with certain microbial taxa and inflammation^9^. After oral surgeries like cleft palate repair, antibiotics are often given but it remains unknown whether antibiotics impact wound healing or microbiome composition after the procedure. Pre-operative assessment of oral surgery patients does not currently mandate microbial assessment or mitigation of associated periodontal disease.

Oral cavity wound healing is less well characterized compared to cutaneous or intestinal wound healing. It is known that there is less scar tissue formation, and that oral wound healing happens faster than cutaneous wound healing^10^, even though oral cavity wound healing occurs in an environment that undergoes constant trauma during eating^11^. Wound healing following cleft palate repair involves separation of the oral and nasal cavities achieved by elevating and rotating the palate tissues to the midline. Many factors that are associated with the occurrence of an oro-nasal fistula (ONF) play a role in healing of these rotated mucosal flaps, including wound tension, width of the cleft palate, and nutritional status^12^. To mitigate the occurrence of ONF formation, surgeons have used multiple strategies to reduce ONF formation by implanting a physical barrier (donated human dermis) to improve wound healing or add an additional layer of the closure to reduce ONF formation. Reports suggest implanting donated human dermis does reduce ONF frequency, but this approach also carries the risk of prion/HIV transmission, and the dermis is only acting as a physical barrier ^13,14^. Reliable local or free tissue flaps of tissue are often a last resort, including the attachment of tongue tissue to the ONF or transfer of a free tissue flap from the radius or the thigh.

Herein, we report that a freshly inflicted wound in the oral palate of a mouse that models an open and unhealed ONF resulted in a significant lowering of oral microbiome alpha diversity, and in blooms of *Enterococcus faecalis, Staphylococcus lentus, and Staphylococcus xylosus* in the oral cavity. Treatment of mice with antibiotics prior to ONF infliction lowered microbiome alpha diversity, prevented *E. faecalis, S. lentus, and S. xylosus* blooms, but did not impact ONF healing. However, delivery of the pro-tissue restitutive probiotic *Lactococcus lactis* subsp. cremoris (LLC) to the ONF wound bed via a PEG-MAL hydrogel led to rapid ONF healing, preserved microbiome alpha diversity, and limited the abundance of *E. faecalis, S. lentus, and S. xylosus* in the oral cavity. Together, we show that a freshly inflicted ONF in the murine palate is associated with a dysbiotic oral microbiome state and ONF formation, and a bloom of opportunistic pathogens. We also show that that delivery of a specific beneficial microbe, LLC, to the ONF can improve wound healing, can restore and/or preserve oral microbiome diversity, and inhibit blooms of opportunistic pathogens.

## Results

### A mouse model of oro-nasal fistula (ONF)

To develop a mouse model of an oro-nasal fistula (ONF), 10-week-old C57BL/6 mice were purchased from Jackson Laboratories and acclimatized at the Emory University mouse vivarium for two weeks. The 12-week old C57BL/6 mice were inflicted with a midline 1.5 mm injury in the hard palate. During 7 days after wound formation, there was gradual narrowing of the wound. The initial injury was associated with exposure of the midline vomer bone, like that in humans, which narrowed over 7 days, as is the case in human healing (**Fig. 1 A-B**). Analysis of the hard palate through the ONF at day 5 revealed detectable tissue re-epithelialization at the edges of the wound (**Fig. 1 C-D**). However, healing did not completely seal the wound, and an opening between the oral cavity and the nasal cavity persisted, as is the case in the formation of an ONF in humans (**Fig. 1**). Thus, this establishes a faithful model for human ONF in mice.

**Fig. 1:**
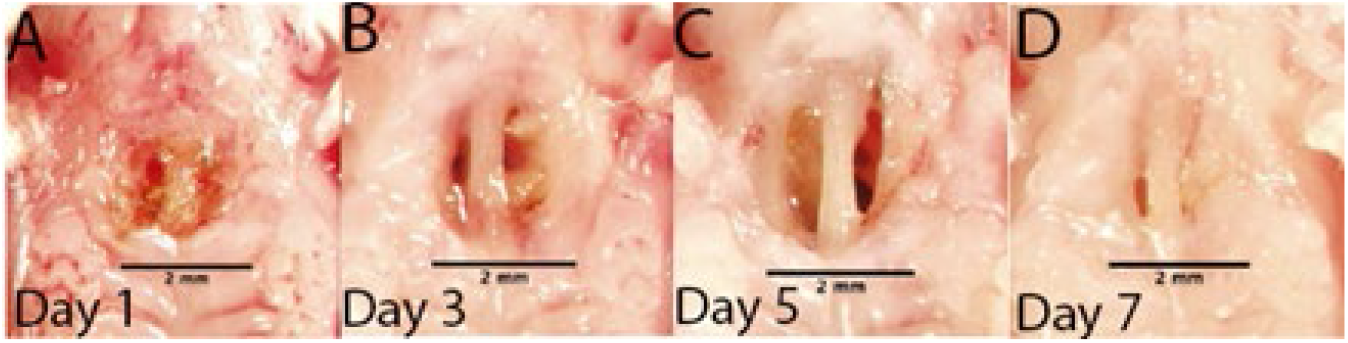
Mouse model of ONF demonstrating the occurrence of the fistula. **(A-B)** day 1 and day 3 after ONF wound infliction. **(C-D)** gradual narrowing of the fistula at 5 days and final ONF formation at day 7

### Oral Microbiome Changes During ONF Wound Healing

Our research group previously reported that the microbiome composition changes following a biopsy wound to colonic tissue^15^. Based on this premise, we characterized the oral microbiome composition in mice before and after ONF injury. We show first that creation of an ONF in mice resulted in significant lowering of oral microbiome alpha diversity, with concurrent blooms of *Enterococcus faecalis, Staphylococcus lentus, and Staphylococcus xylosus* in the oral cavity (**Fig. 2A-C**). We also show by beta diversity analysis that an ONF results in a persistent change in the microbiome composition up to 7 days post injury (**Fig. 2D**). Furthermore, an ONF resulted in increased total bacterial burden in the oral cavity with significantly higher 16S rDNA copy number per microliter sample detected after ONF (**Fig. 2E**). These data outlined a significant change in the Oral Microbiome after ONF formation.

**Figure 2.**
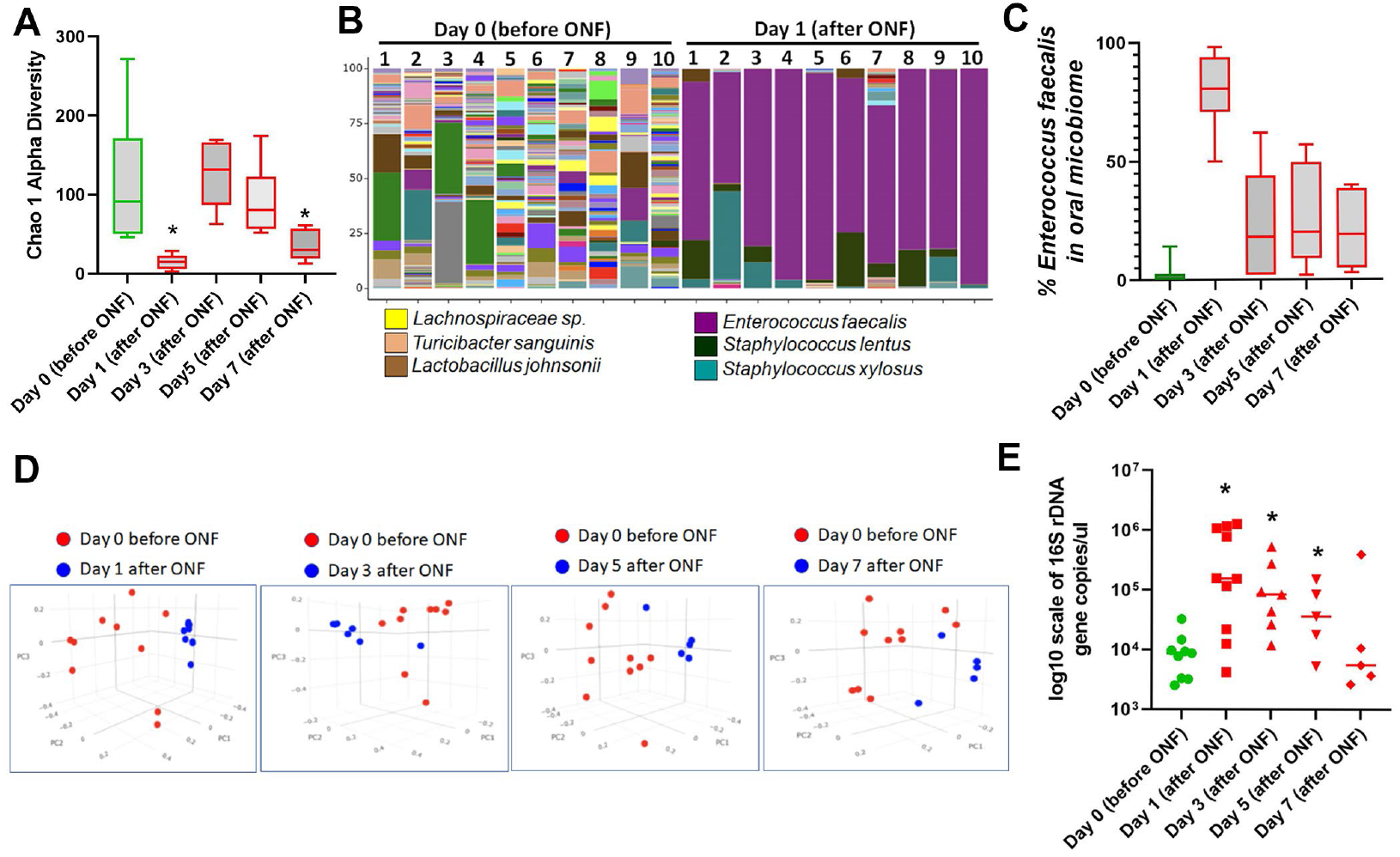
Analysis of the microbiome composition within the oral cavity of ONF wound model mice. **(A)** Measurement of the alpha diversity within oral microbiome samples before (day 0) and after ONF wound infliction for 7 days. Note, the Chao1 alpha diversity measures the microbial diversity of each sample and is a representation of the number of observed species in the samples. **(B)** Histogram representation of the alpha diversity within oral microbiome samples before (day 0) and at day 1 after ONF wound infliction. Note expansion in *Enterococcus faecalis* at day 1. **(C)** Percentage of the total microbiome that is attributed to *Enterococcus faecalis* within the oral microbiome before (day 0) and after ONF wound infliction for 7 days. **(D)** Change in microbiome beta diversity in the oral cavity of mice subjected to an ONF for up to 7 days, compared to the microbiome diversity at day 0 before wound infliction. Beta diversity is a measurement of microbial diversity differences between samples. The figure is the 3-dimensional Principal Coordinate Analysis (PCoA) plot created using the matrix of paired-wise distance between samples calculated by the Bray-Curtis dissimilarity using unique amplicon sequence variants (ASV). Note: ONF results in a persistent change in the microbiome composition over at least 7 days. **(E)** The absolute abundance of bacterial (16S) DNA measured in the oral cavity at day 0 before ONF and after ONF wound infliction for 7 days. Note ONF resulted in increased total bacterial burden in the oral cavity with significantly higher 16S rDNA copy number per microliter detected after ONF. Statistical analysis by t-test with p values shown on graphs. *=P=<0.05.

### Antibiotic Treatment impacts the Oral Microbiome During ONF Wound Healing

*E. faecalis* is associated with oral diseases, such as caries, endodontic infections and periodontitis^16,17^. However, the implications of these changes to the oral microbiome composition, both loss of taxa diversity or alpha diversity, and the bloom of other taxa such as *E. faecalis*, to tissue healing processes remains unstudied. Moreover, whether interactions of specific oral microbes with host cells activate pro-restitutive cellular signaling pathways in the oral wound bed is also unknown. To this end, we treated mice with a cocktail of 1mg/mL Ampicillin, 0.5 mg/mL Vancomycin, 1 mg/mL Neomycin, 1 mg/mL Metronidazole included in their drinking water, which are absorbable and non-absorbable antibiotics commonly used in experimental science to deplete the murine microbiome of normal viability^18^, for 1 week and detected a > 99.9% lowering in total bacterial burden (**Fig. 3A**). We also detected an expected reduction in the microbiome alpha diversity (**Fig. 3B**). Antibiotic treatment rendered *E. faecalis* undetectable in the oral microbiome at day 0 (before ONF), and throughout the 7 days over which the rate of healing is followed (**Fig. 3C**). However, the antibiotic-mediated elimination of the bacterial load in the oral cavity but did not impact ONF healing, with ONF still visible in the palate of mice at day 7 after wound infliction (**Fig. 3D**). These data suggest that antibiotics impact oral microbiome composition, but do not promote wound healing. The data collectively suggest that a dysbiotic microbiome (**Fig. 2**), or loss of microbiome diversity (**Fig. 3**) may both impair wound healing and may contribute to the formation of an ONF.

**Figure 3.**
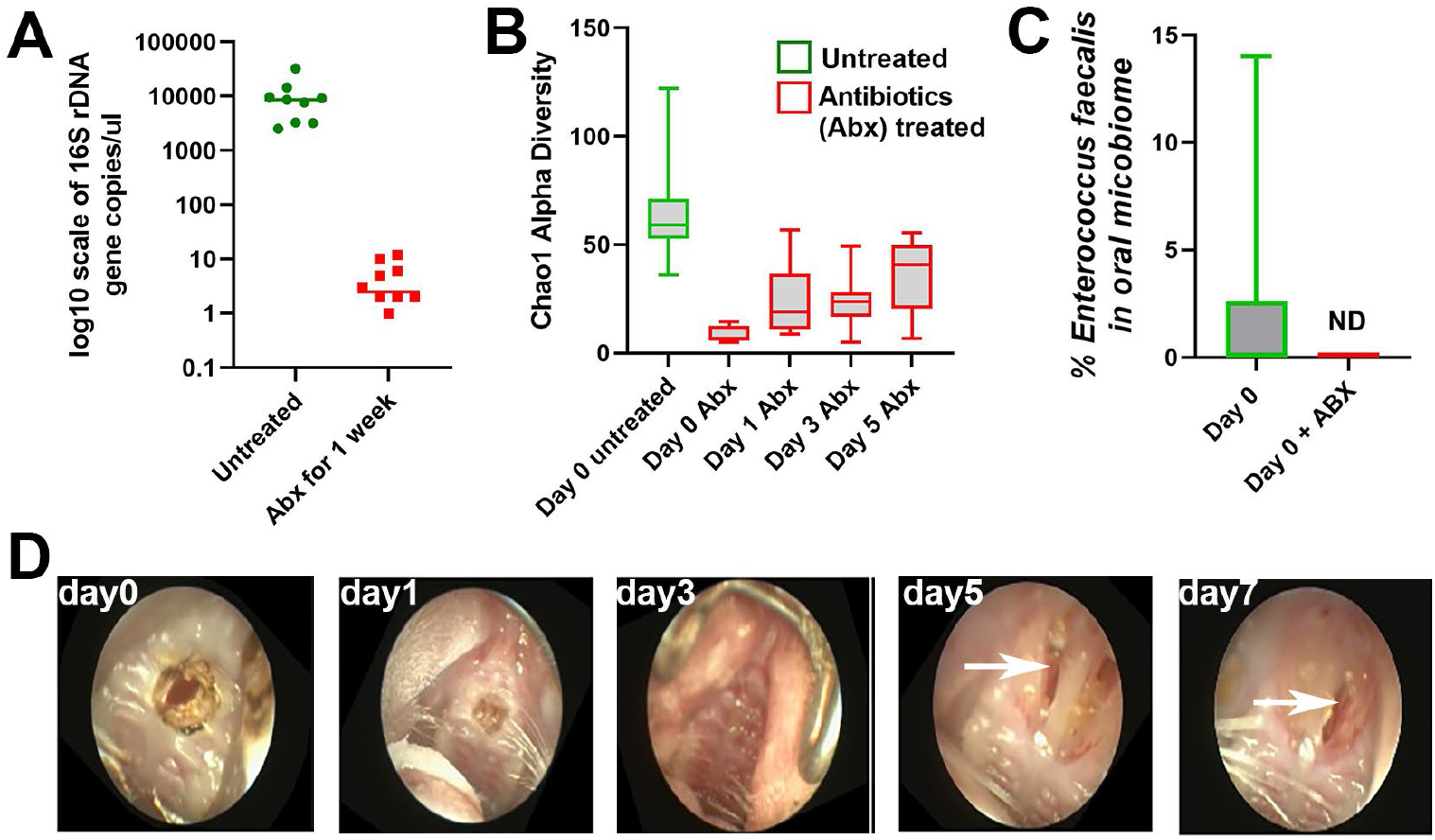
Antibiotic Treatment impacts the Oral Microbiome During ONF Wound Healing. **(A)** The absolute abundance of bacterial (16S) DNA measured in the oral cavity mice treated with of antibiotics (Ampicillin, Vancomycin, Neomycin, Metronidazole) for 7 days, compared to untreated mice. **(B)** Chao1 alpha diversity within the oral microbiome of mice treated with antibiotic as described in (F) and subjected to an ONF as described in figure 2. **(C)** Percentage of the total microbiome that is attributed to *Enterococcus faecalis* within the oral microbiome mice treated with antibiotic as described in (F) at day 0 before ONF, compared to untreated controls. ND=not detected. **(D)** Wound healing of ONF in mice treated with antibiotics as described in (F), for up to 7 days post wound infliction. Note. ONF is still visible (white arrows) in mice treated with antibiotics mice after 7 days. Images are a representative of 10 biological replicates. Experimental power: For mice in A to E, day 0 (n=10), day 1 (n=8), day 3 (n=6), day 5 and 7 (n=5). For mice in A to D (antibiotic experiment) day 0 to day 7 (n=10). Statistical analysis by t-test with p values shown on graphs. *=P=<0.05.

### Beneficial Microbes enhance ONF Wound Healing

An alternative strategy to promote tissue wound healing and prevent the formation of an ONF is the use of beneficial microbes, commonly known as ‘probiotics’, which are defined as, ‘live microorganisms which when administered in adequate amounts confer a health benefit on the host.’ Current challenges in probiotic usage involve the development of individualized microbial therapeutics for targeted intervention, an approach defined by as “Precision probiotic therapy”. This involves selecting probiotics with known specific activates, such as in the healing of an ONF, and the activation of pro-restitutive tissue healing pathways. A further challenge in the context of oral wound healing is that the wound in an environment that undergoes constant trauma during eating. In response to these challenges, we developed an approach where we selected a pro-tissue restitutive probiotic, namely *Lactococcus lactis* subsp. cremoris (LLC)^19,20^, that we deliver directly into the ONF wound in the oral palate using a polyethylene glycol malamide (PEG-MAL) hydrogel (**Fig 4A**). Strikingly, delivery of the pro-tissue restitutive probiotic *L. lactis* subsp. cremoris (LLC) to the wound bed of the freshly inflicted ONF via a PEG-MAL hydrogel vehicle resulted in rapid healing of the ONF (**Fig 4B-C**). Healing of the inflicted wound and the absence of an ONF was associated with the preservation of relatively high microbiome alpha diversity and limited the abundance of *E. faecalis* in the oral cavity (**Fig 4D-F**). The data demonstrate that delivery of a specific beneficial microbe, LLC, to the ONF can boost wound healing, can restore and/or preserve oral microbiome diversity, and inhibit blooms of opportunistic pathogens.

**Figure 4.**
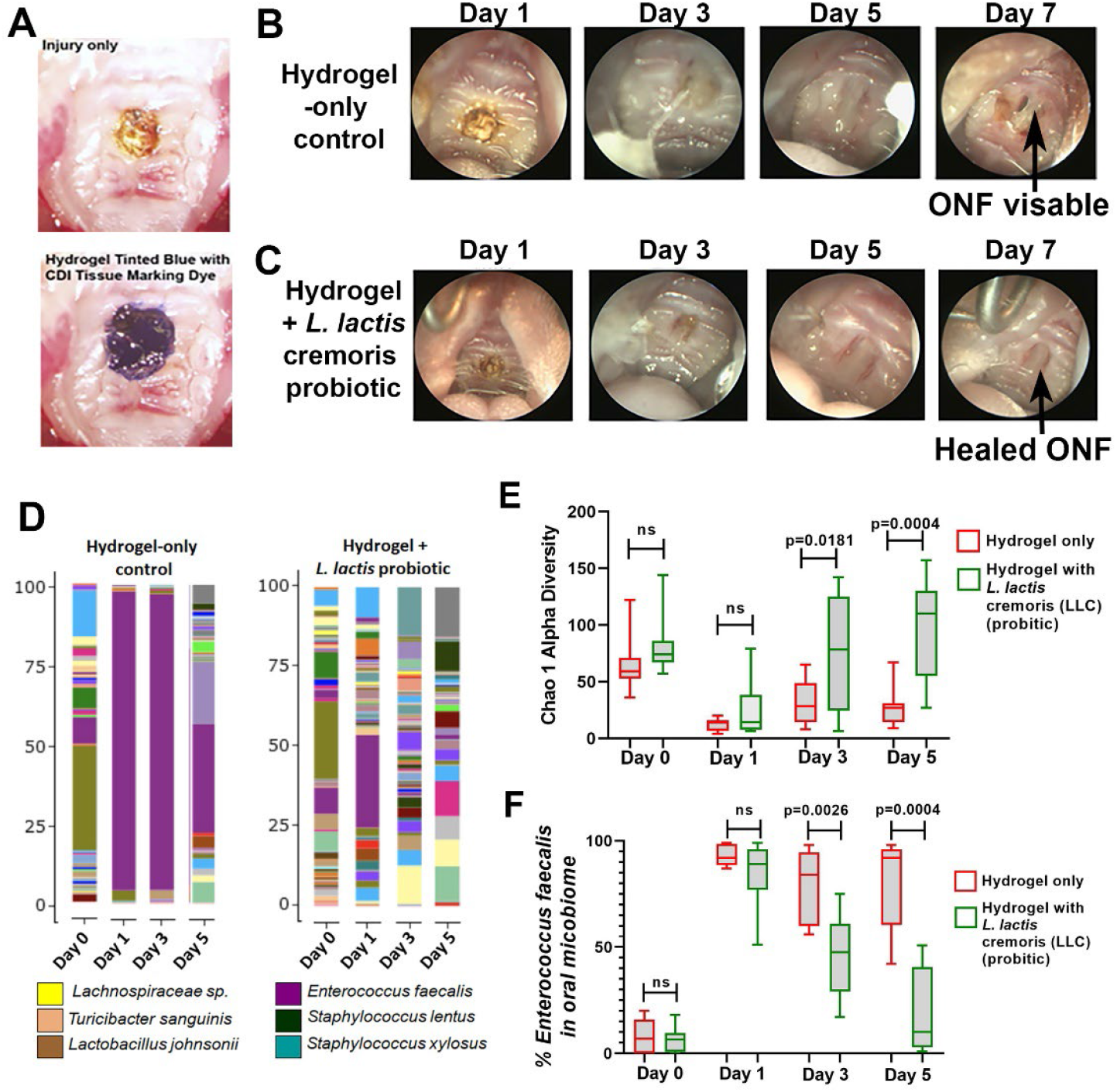
Delivery of a probiotic to the ONF wound site using hydrogel technology promotes ONF wound healing. **(A)** As proof of principle, we show our expertise in applying hydrogel to the site of an oral ONF wound formed as described in figure 2. We also tinted the hydrogel with a blue CDI tissue marking dye for enhanced observation. We detected that the hydrogel remains within the wound for 24 hours following application. In out experiments, the hydrogel containing probiotic is reapplied every 24 hours. **(B and C)** Representative images of mice inflicted with ONF as described in figure 2, and then treated daily with polyethylene glycol malamide (PEG-MAL) hydrogel only control (panel B), or a PEG-MAL) hydrogel containing 1×10E6 CFU *Lactococcus lactis* subspecies cremoris (LLC) (panel C). Note ONF is still visible (black arrows) in hydrogel only control (panel B) mice after 7 days, but ONF is healed (black arrows) in mice treated with hydrogel containing 1×10E6 CFU LLC (panel C) after 7 days. **(D)** Histogram representation of the alpha diversity within oral microbiome of samples before (day 0) and for up to 5 days after ONF wound infliction in a representative median mouse from each group described in B and C. Note expansion of *Enterococcus faecalis* in hydrogel only control mice, but markedly less *E. faecalis* detected in hydrogel containing LLC treated mice. **(E)** Measurement of the Chao1 alpha diversity within oral microbiome samples of mice described in B and C above. Note significantly higher alpha diversity in mice treated with hydrogel containing LLC compared to control. **(F)** Percentage of the total microbiome that is attributed to *Enterococcus faecalis* within the oral microbiome of samples described in B and C above. Note significantly lower amounts of *E. faecalis* in mice treated with hydrogel containing LLC compared to control. In all experiments, n=10 mice. Statistical analysis by t-test with p values shown on graphs.

## Discussion

Cleft palate formation is the most common congenital craniofacial anomaly (1:700 births). Adverse healing after cleft palate repair occurs in up to 60% of children and often leads to an oro-nasal fistula (ONF), which is a direct opening between the mouth and the nose that requires multiple surgeries to repair. Healing in the oral cavity occurs in the setting of constant physical trauma and a milieu of bacterial, fungal, parasite and viral organisms. ONF is a serious impediment to a child’s ability to eat and talk. Therefore, there is an urgent need to understand the microbiological context in which ONF healing occurs and how the oral microbiome can be optimized to improve ONF healing. Development of novel therapeutic strategies to enhance healing after cleft palate surgery to reduce the occurrence of ONF would reduce morbidity, improve outcomes, and reduce costs.

It has been shown that the microbiota in the gut elicit profound effects on host gene regulatory events within surgical wounds^21,22^. In addition, it is well established that wound healing on the skin and in the intestine is sensitive to local environmental factors, including the microbiome, where certain microbial community structures or probiotic bacteria can impede or enhance the wound healing process respectively. We contribute to a new body of knowledge about how an eubiotic or dysbiotic oral microbiome affects oral wound healing. We determined that antibiotics during oral surgery lowers the microbial burden in the oral cavity but does not enhance wound healing of an ONF. Strikingly, delivery of a pro-regenerative beneficial microbe, LLC into the ONF resulted in complete healing of the ONF and restored the microbial diversity. Overall, we show that restoring an eubiotic oral microbiome exerts positive modulatory influences on oral wound healing, plausibly through bacterial-bolstered immune and epithelial cell regenerative cues. Discovering specific bacterial taxa within the oral cavity or ONF that positively influence healing will be essential information to characterize the overall constituants of eubiotic oral microbiome.

Together, these data will yield critical data to inform new therapeutic modalities to lower the prevalence of ONFs following cleft palate surgery and may be expanded to other clinical use cases related to oral wound healing following trauma and cancer surgery.

Mechanistically, a critical step of tissue repair occurs during the inflammatory stage, and this is contingent on the mobilization of circulating monocytes into the wound bed^23^. Based on local factors, these monocytes undergo differentiation into inflammatory (IM) and antiinflammatory (AM) monocytes^24^. It’s also known that IM and AM are exquisitely sensitive to the quantity and quality of the oral microbiome composition. Relevant to ONF healing is that notion that a dysbiotic oral microbiome detected following a ONF infliction that is enriched in *E. faecalis, S. lentus, and S. xylosus* may provoke an aggravated immune response that impedes tissue healing.

On the other hand, elimination of commensal background microbiome using a broad-spectrum antibiotic cocktail, although eliminating *E. faecalis, S. lentus, and S. xylosus*, did not improve ONF healing. Indeed, these data suggest that a near complete absence of bacteria (99.9% less bacterial load than untreated), and blunting of an immune cell response does not result in a pro-restitutive wound healing events in the oral tissue. The discovery that delivery of specific strain of probiotic bacteria, i.e. LLC directly into the ONF via a hydrogel can promotes ONF healing opens a window of opportunity to examine the specific gene regulatory events within an oral palate that is characteristic of a pro-restitutive environment. Discovering these mechanisms will be a challenging future endeavor.

## Material and Methods

### Experimental Mice

All experiments were done using C57BL/6 mice purchased from The Jackson Laboratories. Animal procedures were approved by the Institutional Animal Care and Use Committee of Emory University.

### Mouse ONF Model

Using 12-week-old C57BL/6, mice, we provided anesthesia using ketamine and buprenorphine to perform the surgery. The mice were placed in a supine position and a dissecting microscope is visualize the oral cavity. An eye lid retractor was used to retract the upper and lower jaw with the tongue pushed out of the way. Using an ophthalmologic cautery, a 1.5 mm oronasal fistula was performed between the first molars in the midline and hemostasis was achieved. In the control mice, the defect alone was created. In the hydrogel mice, the hydrogel (described below) was placed into the wound after gel formation had occurred and a Tegaderm bandage was placed over the hydrogel and cyanoacrylate adhesive was used to fix the Tegaderm bandage to the surrounding oral tissues.

### Analysis of Microbiome sequencing 16S rDNA sequencing

Oral microbiome samples were collected from 12-week-old mice using swabs. The swabs were analyzed through sequencing of the 16S rDNA gene undertaken by ZymoBIOMICS (Zymo Research, Irvine, CA). DNA was extracted from swabs using ZymoBIOMICS-96 MagBead DNA kit (Zymo Research, Irvine, CA). The ZymoBIOMICS Microbial Community Standard was used as a positive control for each DNA extraction. The ZymoBIOMICS Microbial Community DNA Standard was used as a positive control for each targeted library preparation. Negative controls were included to assess the level of bioburden carried by the wet-lab process. The DNA samples were then prepared for targeted sequencing with *Quick*-16S Primer Set V3-V4 Library (Zymo Research, Irvine, CA). The sequencing library was prepared using innovative library preparation process in which PCR reactions were performed in real-time PCR machines to control cycles and limit PCR chimera formation. The final PCR products were quantified with qPCR fluorescence readings and pooled together based on equal molarity. The final pooled library was cleaned up with the Select-a-Size DNA Clean & Concentrator (Zymo Research, Irvine, CA), then quantified with TapeStation (Agilent Technologies, Santa Clara, CA) and Qubit (Thermo Fisher Scientific, Waltham, WA). The final library was sequenced on Illumina MiSeq with a v3 reagent kit (600 cycles). Calculations of alpha and beta diversity were done by standard methodology by Zymo Research.

### Bacterial strains and culture preparation and administration of Bacteria

The following bacteria were purchased from the American Type Culture Collection (ATCC) (Manasas, VA): *Lactococcus lactis* subspecies cremoris ATCC 19257. All media was propagated according to instructions provided by the ATCC.

### Creation and delivery of PEG-MAL hydrogel with *Lactococcus lactis* subsp. cremoris (LLC)

To produce the hydrogel, PEG-MAL hydrogel a twenty-kilodalton PEG-4MAL macromer (Lysan Bio) was cross-linked in a one-step reaction by combining PEG-lysostaphin with the GFOGER peptide, GGYGGP(GPP)5GFOGER(GPP)5GPC (New England Peptide) and the bacterial suspension. Bacterial suspensions were prepared by picking individual colonies of bacteria grown on a TSA plate overnight and suspending them in Dulbecco’s PBS supplemented with calcium and magnesium (PBS) to an optical density of 0.20 at 600 nm (MicroScan Turbidity Meter; Seimens) and then diluting this suspension 100-fold. The viable count for all bacterial inocula was determined by plate count on the relevant culture medium. The hydrogel was 4.0% wt/vol 20-kDa PEG-4MAL, 1 mM GFOGER, and 424 U/mL of LLC. The amount of VPM cross-linker added was determined stoichiometrically by matching the remaining maleimide groups after accounting for GFOGER incorporation. After mixing, the hydrogels were allowed to gel for 15 min in a humidified incubator at 37 °C and 5.0% CO2 for implantation into the ONF.

### Statistical Analysis

One-way ANOVA, with Bonferroni’s multiple comparisons test, was used for analysis of data. Wound healing assay assay data were analyzed using a Kruskal-Wallis test with Dunn’s multiple comparisons. Two-way ANOVA with Bonferroni’s multiple comparisons was used for analysis of proliferation data. Differences were considered statistically significant for P-values < 0.05. All analyses were performed using GraphPad Prism v6.0.4 software (GraphPad, San Diego, CA, USA). Data are presented as the mean ± standard error of the mean (SEM).

## Author Contributions

SLG and RMJ conceived and designed the experiments. SLG, HB and AT performed experiments for ONF wound infliction. CRN and CAG performed analyzed 16S rDNA sequencing. WW, MT, VS and AC prepared hydrogel formulation for delivery of probiotics to ONF. l. SLG and RMJ wrote the manuscript.

## Acknowledgments

This research was funded by the National Institutes of Health, National Institute of Dental and Craniofacial Research to SLG and RMJ (R21 DE030632). RMJ is supported, in part, by a startup fund from The Emory University School of Medicine.

## References

1. Deo, P.N. & Deshmukh, R. Oral microbiome: Unveiling the fundamentals. J Oral Maxillofac Pathol 23, 122–128 (2019).

2. Sharma, N., Bhatia, S., Sodhi, A.S. & Batra, N. Oral microbiome and health. AIMS Microbiol 4, 42–66 (2018).

3. Dong, L., Yin, J., Zhao, J., Ma, S.R., Wang, H.R., Wang, M., Chen, W. & Wei, W.Q. Microbial Similarity and Preference for Specific Sites in Healthy Oral Cavity and Esophagus. Front Microbiol 9, 1603 (2018).

4. Willis, J.R. & Gabaldon, T. The Human Oral Microbiome in Health and Disease: From Sequences to Ecosystems. Microorganisms 8(2020).

5. Gomez, A., Espinoza, J.L., Harkins, D.M., Leong, P., Saffery, R., Bockmann, M., Torralba, M., Kuelbs, C., Kodukula, R., Inman, J., Hughes, T., Craig, J.M., Highlander, S.K., Jones, M.B., Dupont, C.L. & Nelson, K.E. Host Genetic Control of the Oral Microbiome in Health and Disease. Cell Host Microbe 22, 269–278 e3 (2017).

6. Gomez, A. & Nelson, K.E. The Oral Microbiome of Children: Development, Disease, and Implications Beyond Oral Health. Microb Ecol 73, 492–503 (2017).

7. Dewhirst, F.E. The Oral Microbiome: Critical for Understanding Oral Health and Disease.J Calif Dent Assoc 44, 409–10 (2016).

8. Wade, W.G. The oral microbiome in health and disease. Pharmacol Res 69, 137–43 (2013).

9. Liu, L., Zhang, Q., Lin, J., Ma, L., Zhou, Z., He, X., Jia, Y. & Chen, F. Investigating Oral Microbiome Profiles in Children with Cle Lip and Palate for Prognosis of Alveolar Bone Graing. PLoS One 11, e0155683 (2016).

10. Mak, K., Manji, A., Gallant-Behm, C., Wiebe, C., Hart, D.A., Larjava, H. & Hakkinen, L. Scarless healing of oral mucosa is characterized by faster resoluon of inflammaon and control of myofibroblast action compared to skin wounds in the red Duroc pig model. J Dermatol Sci 56, 168–80 (2009).

11. Chen, L., Arbieva, Z.H., Guo, S., Marucha, P.T., Mustoe, T.A. & DiPietro, L.A. Positional differences in the wound transcriptome of skin and oral mucosa. BMC Genomics 11, 471 (2010).

12. Phua, Y.S. & de Chalain, T. Incidence of oronasal fistulae and velopharyngeal insufficiency aer cle palate repair: an audit of 211 children born between 1990 and 2004. Cleft Palate Craniofac J 45, 172–8 (2008).

13. Kirschner, R.E., Cabiling, D.S., Slemp, A.E., Siddiqi, F., LaRossa, D.D. & Losee, J.E. Repair of oronasal fistulae with acellular dermal matrices. Plast Reconstr Surg 118, 1431–40 (2006).

14. Gonzalez-Sanchez, J.G. & Jimenez-Barragan, K. [Closure of recurrent cle palate fistulas with plasma rich in growth factors]. Acta Otorrinolaringol Esp 62, 448–53 (2011).

15. Alam, A., Leoni, G., Quiros, M., Wu, H., Desai, C., Nishio, H., Jones, R.M., Nusrat, A. & Neish, A.S. The microenvironment of injured murine gut elicits a local pro-restitutive microbiota. Nat Microbiol 1, 15021 (2016).

16. Kouidhi, B., Zmantar, T., Mahdouani, K., Hentati, H. & Bakhrouf, A. Anbioc resistance and adhesion properes of oral Enterococci associated to dental caries. BMC Microbiol 11, 155 (2011).

17. Rams, T.E., Feik, D., Mortensen, J.E., Degener, J.E. & van Winkelhoff, A.J. Anbioc suscepbility of periodontal Enterococcus faecalis. J Periodontol 84, 1026–33 (2013).

18. Tyagi, A.M., Yu, M., Darby, T.M., Vaccaro, C., Li, J.Y., Owens, J.A., Hsu, E., Adams, J., Weitzmann, M.N., Jones, R.M. & Pacifici, R. The Microbial Metabolite Butyrate Smulates Bone Formaon via T Regulatory Cell-Mediated Regulation of WNT10B Expression. Immunity 49, 1116–1131 e7 (2018).

19. Darby, T.M., Naudin, C.R., Luo, L. & Jones, R.M. Lactobacillus rhamnosus GG-induced Expression of Lepn in the Intesne Orchestrates Epithelial Cell Proliferaon. Cell Mol Gastroenterol Hepatol 9, 627–639 (2020).

20. Naudin CR O.J., Askew LC, Khan RN, Scharer CD, Mathews JD, Luo L, Kim J, Reedy AR, Barbian ME, and RM Jones. Lactococcus lactis sb. cremoris orchestrates signal events in the gut epithelium via TLR2 to promote ssue restuon. BioRxiv 471025 [Preprint](2021).

21. Alam, A., Leoni, G., Quiros, M., Wu, H.X., Desai, C., Nishio, H., Jones, R.M., Nusrat, A. & Neish, A.S. The microenvironment of injured murine gut elicits a local pro-restitutive microbiota. Nature Microbiology 1(2016).

22. Alam, A., Leoni, G., Wentworth, C.C., Kwal, J.M., Wu, H., Ardita, C.S., Swanson, P.A., Lambeth, J.D., Jones, R.M., Nusrat, A. & Neish, A.S. Redox signaling regulates commensal-mediated mucosal homeostasis and restuon and requires formyl pepde receptor 1. Mucosal Immunology 7, 645–655 (2014).

23. Mathews, J.D., Owens, J.A., Naudin, C.R., Saeedi, B.J., Alam, A., Reedy, A.R., Hinrichs, B.H., Sumagin, R., Neish, A.S. & Jones, R.M. Neutrophil-Derived Reactive Oxygen Orchestrates Epithelial Cell Signaling Events during Intestinal Repair. Am J Pathol 189, 2221–2232 (2019).

24. Klinkert, K., Whelan, D., Clover, A.J.P., Leblond, A.L., Kumar, A.H.S. & Caplice, N.M. Selecve M2 Macrophage Depleon Leads to Prolonged Inflammaon in Surgical Wounds. Eur Surg Res 58, 109–120 (2017).

